## Supplementary material for "A genetic circuit on a single DNA molecule as autonomous dissipative nanodevice": SI

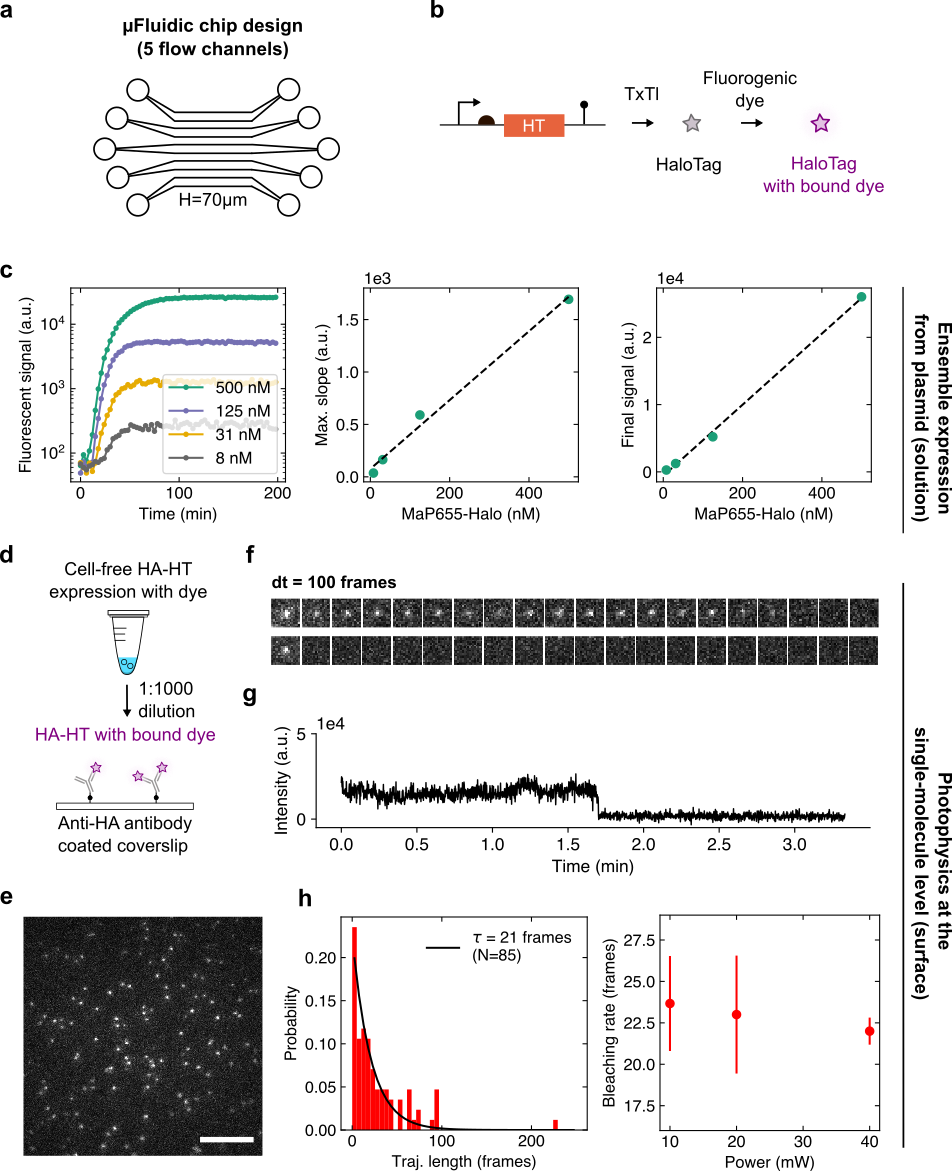

**Supplementary Figure 1: Cell-free expression and photo-physics of MaP655-Halo bound to HT proteins. a)** Microfluidic chip design with 5 parallel flow channels (L~6000 µm, W~600 µm). **b)** Gene expression from the *ht* cassette produces HT proteins that can rapidly bind the fluorogenic dye MaP655-Halo. The fluorogenic dye reacts with nascent HT proteins and increases its fluorescent signal by ~1,000-fold. **c)** Cell-free expression dynamics with various concentrations of MaP655-Halo. As expected, the maximal slope and final fluorescence signal of the expression curves (as shown in the leftmost panel) increased linearly with the concentration of MaP655-Halo. **d)** To measure the photo-stability of bound MaP655-Halo, HA-tagged HT was expressed in solution with 50 nM of fluorogenic dye. The DNA was expressed for ~1 h to produce enough HT to bind most free fluorogenic dye. The mix was diluted by 1,000-fold with PBS and flushed on an anti-HA antibody coated coverslip. **e)** Fluorescence microscopy image of immobilized HT spots. Scale bar, 10 µm. **f)** Two exemplary HT spots over time with characteristic single-molecule bleaching steps. **g)** The intensity trace of a single HT spot over time. **h)** Bleaching rate of the dye bound to HT and fit to mono-exponential decay (black line) at an excitation power of 15 W cm^-2^ (equals an input power of 10 mW). The bleaching rate for different input powers. The error bars are the SD of fitted rates from mono-exponential decay curves (n=3).

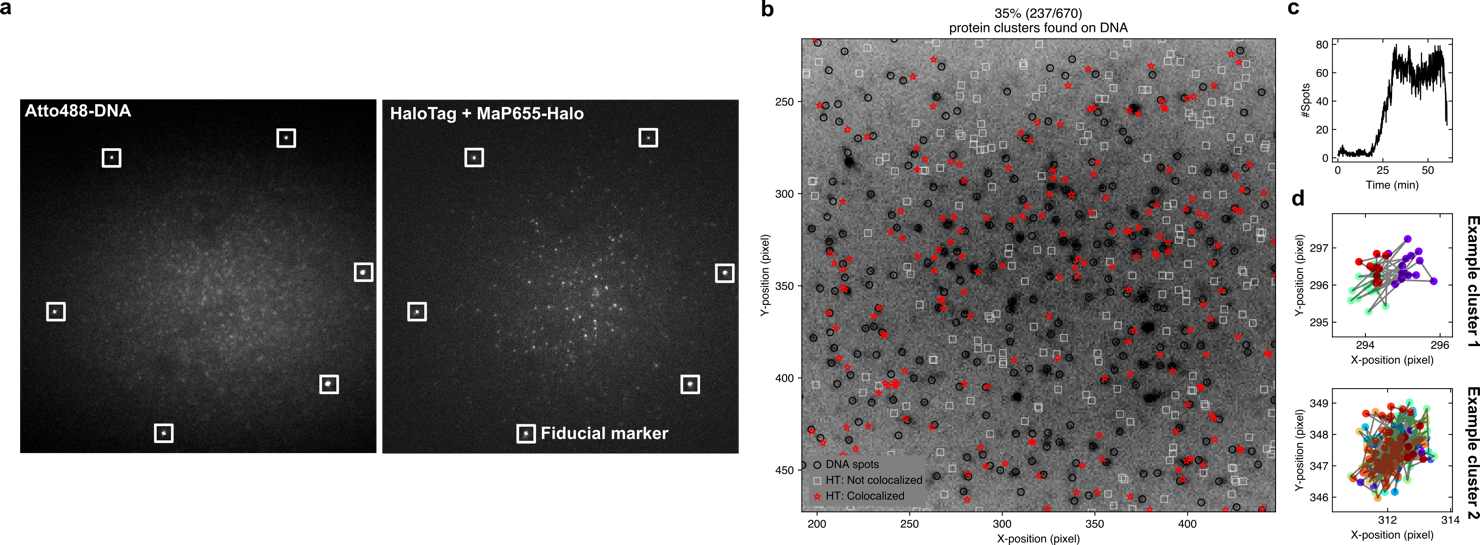

**Supplementary Figure 2: The detection and analysis of protein synthesis spots on the surface of a flow channel. a)** Fluorescent image of DNA molecules and HT protein synthesis spots with fiducial markers (white boxes). **b)** Fluorescence microscopy image of DNA distribution at the beginning of an experiment. The identified DNA spots are indicated by black circles. Colocalized and non-colocalized HT spots are indicated by red stars and white squares, respectively. **c)** The number of detected HT spots over time. The start of expression is marked by an increase of protein synthesis spots on the surface. After tens of minutes, a balance between protein production and removal is reached, maintaining a steady-state protein level on the coverslip surface. **d)** Two examples for identified protein synthesis clusters. The colors indicate protein synthesis spots (one color corresponds to one continuous line of protein synthesis spots) interspersed by periods without protein synthesis spots at the specified location. The individual trajectories are clustered together to define a protein synthesis spot. The average positions for each cluster are used to compute the protein synthesis intensity traces.

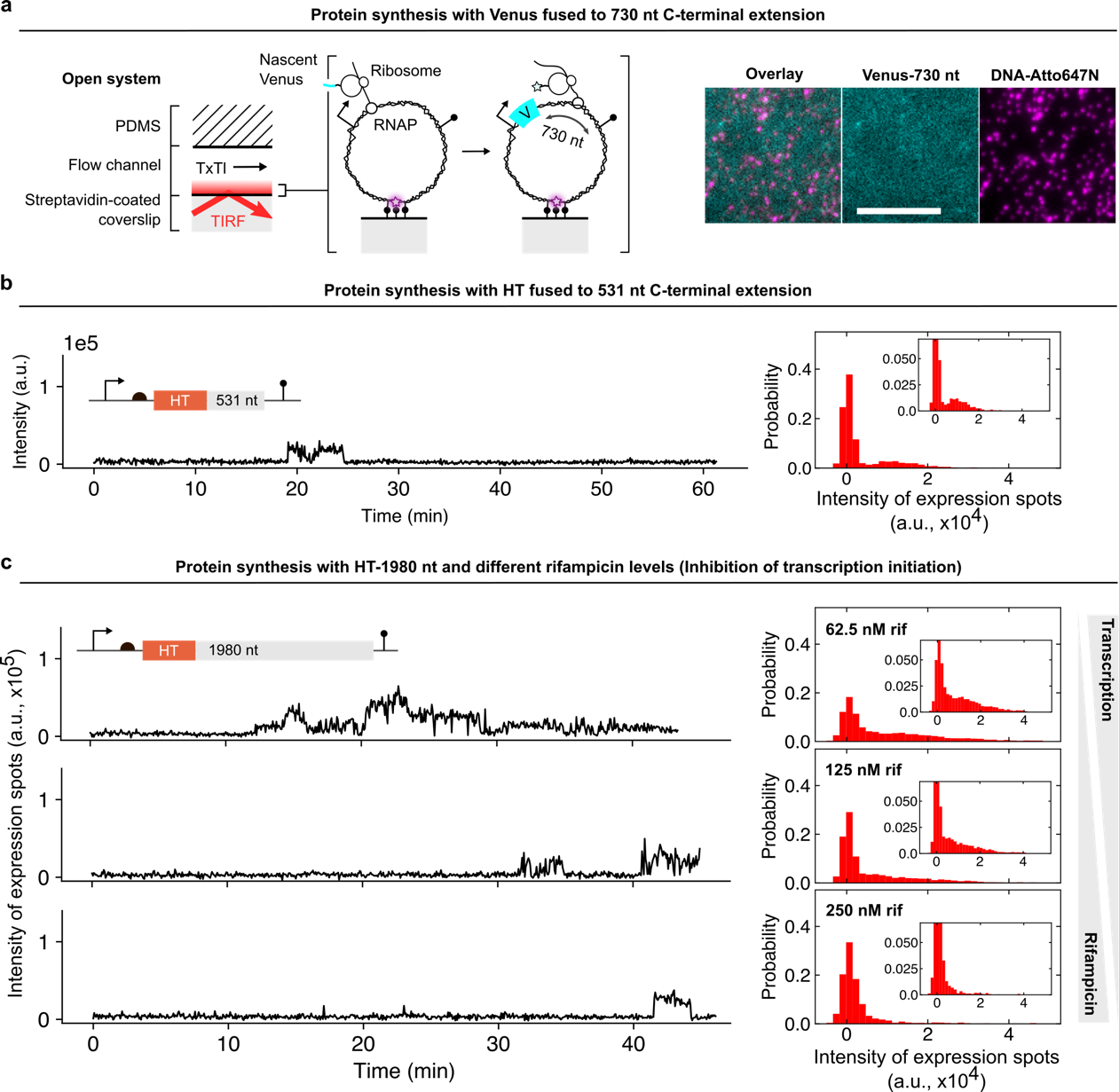

**Supplementary Figure 3: Single-molecule gene expression from DNA molecules encoding Venus, HT fused to a 531 nt long C-extension, and HT fused to 1,980 nt long C-terminal extension with various rifampicin concentrations. a)** Single-molecule gene expression experiment with Venus fused to a 730 nt long C-terminal fusion encoded on Atto647N fluorescently labeled DNA. Scale bar, 10 µm. **b)** Exemplary intensity trace from a single DNA molecule and intensity probability distribution from an ensemble of DNA molecules encoding the *ht* gene fused to a 531 nt-long C-terminal extension. **c)** Exemplary intensity traces for protein synthesis spots (left column) with the 1,980 nt long C-terminal fusion to HT at three rifampicin concentrations as indicated in the intensity probability distributions assembled from an ensemble of DNA molecules (right column).

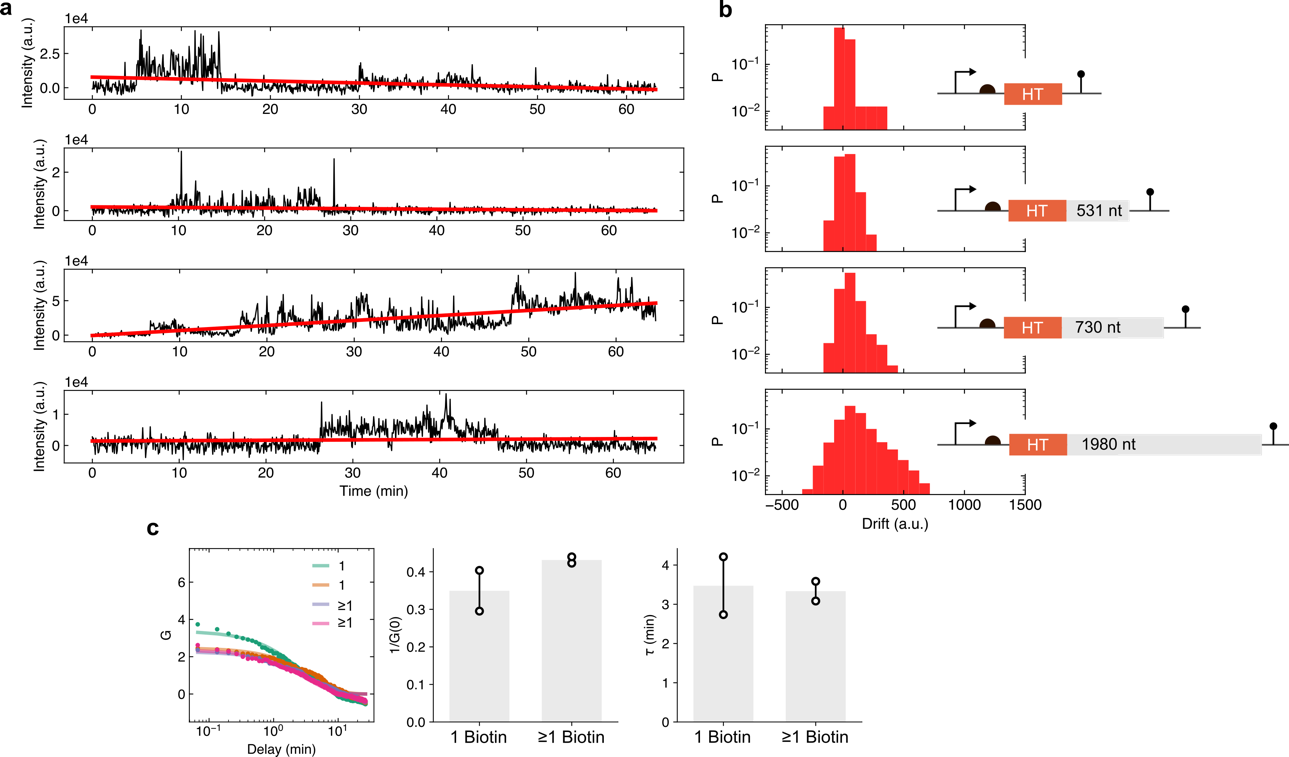

**Supplementary Figure 4: Accumulation of protein signal with different C-terminal extension attached to the HT protein. a)** Exemplary intensity traces for protein synthesis spots with the 2,021 nt long C-terminal fusion and their linear fits (red line). **b)** The slopes of all gene expression spot extracted from the linear fits, plotted as histograms for the four different gene constructs as schematically indicated. **c)** The autocorrelation function, inverse amplitude, and typical time scale as described in the methods and main text for the 2,021 nt long C-terminal fusion length and different numbers of biotins for surface immobilization (n=2).

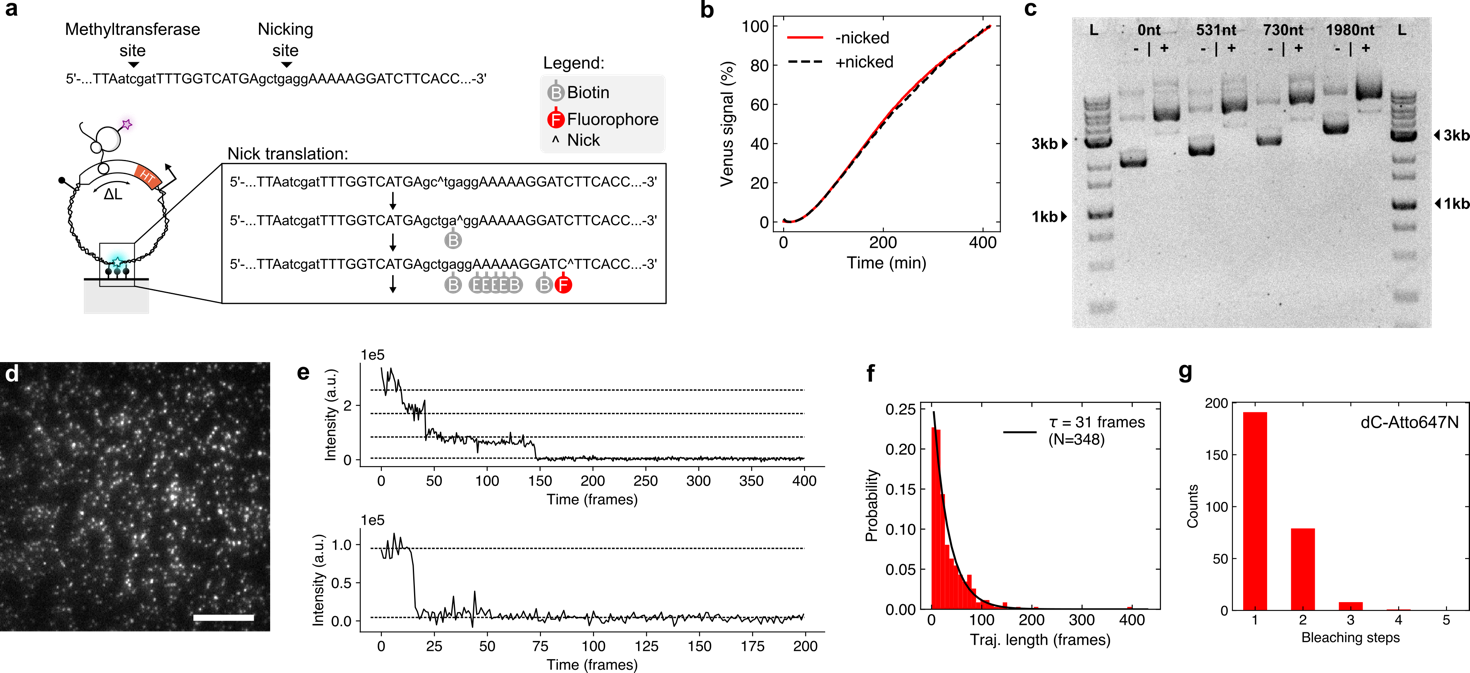

**Supplementary Figure 5: DNA labeling density. a)** Schematic of DNA molecule for surface-immobilization with zoom on sequence containing the two unique sites for the methyltransferase enzyme (M.BseCI) for biotinylating DNA molecules with a single biotin and nicking enzyme (Nb.BbvCI) for modifications with multiple biotins and fluorescent labels (Sequential steps during nick translation are explained in the black box). **b)** Cell-free gene expression in solution from DNA with and without nick. **c)** Agarose gel with DNA encoding the *ht* gene with the four C-terminal fusion tags. The lanes correspond to DNA without (-) and with (+) nicking enzyme after nick translation. The DNA was labeled with Ethidium Bromide and run with a DNA ladder (L). **d)** Image of Atto647N-labeled DNA molecules immobilized on the surface during gene expression. Scale bar, 10 µm. **e)** Typical bleaching trajectories of single DNA. As expected for single dyes, the intensity during imaging exhibited discrete bleaching steps (dashed lines). **f)** The trajectories were extracted and their time until completely bleached was plotted as histogram. The lifetimes were fitted to a mono-exponential decay and gave a typical fluorescent lifetime (=1/bleaching rate) of 31 frames with the excitation parameters that was used in all single-molecule experiments. **g)** Estimated number of fluorophores (by counting bleaching steps) on single DNA molecules.

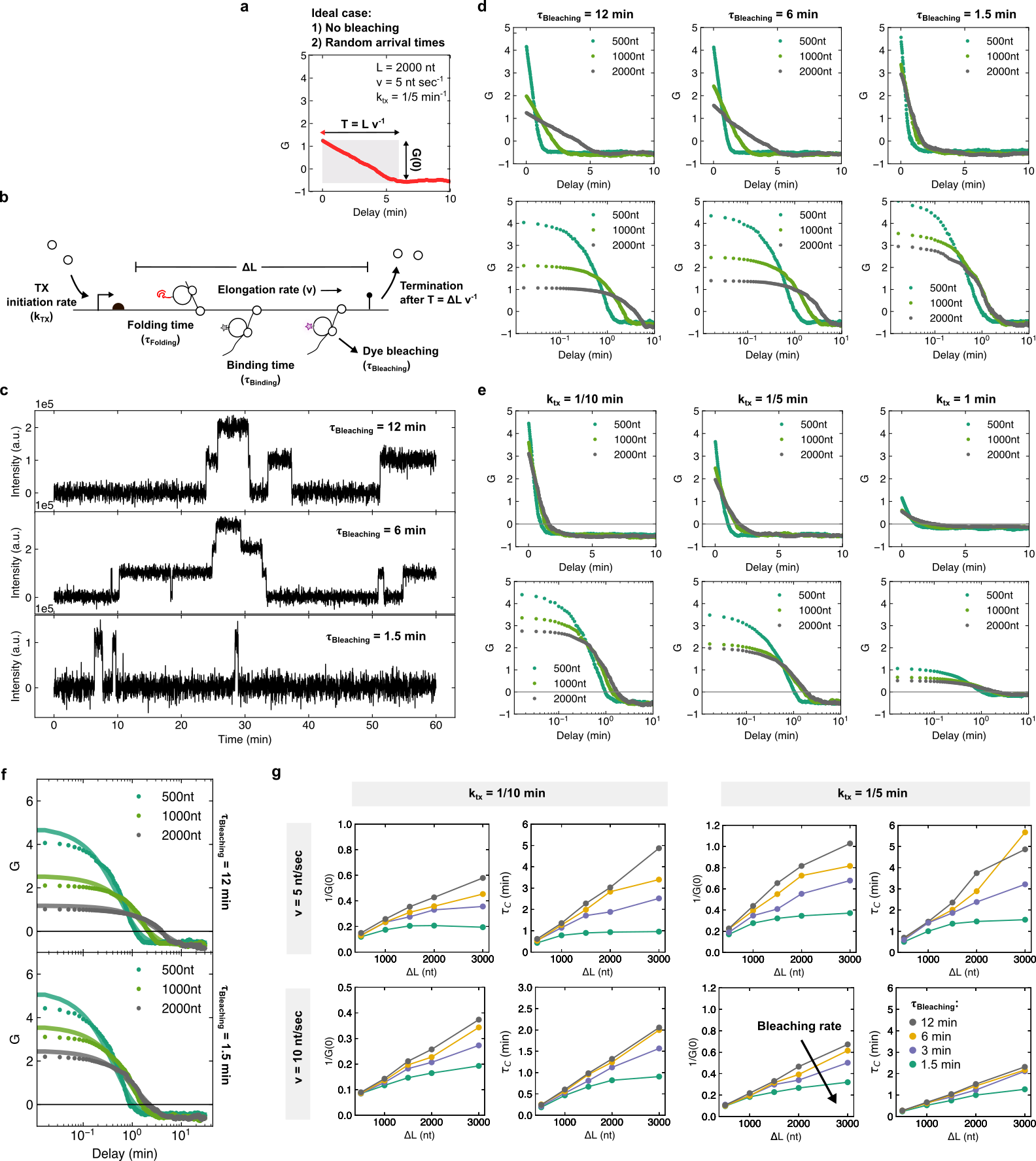

**Supplementary Figure 6: Simulations of ideal and experimentally limited ACF for proteins synthesized through coupled gene expression machines on single DNA. a)** The interpretation of the ideal ACF for a simple model of nascent proteins on single DNA. **b)** Model for dynamics of nascent proteins on DNA. Random transcription initiation events with a rate *k_TX_* occur that transition into transcription and translation complexes moving along the gene with length *L* and velocity *v*. After complete folding of the HT (τ_Folding_), the fluorogenic dye can bind to the nascent HT (τ_Binding_). The fluorescent signal is then either lost due to bleaching (τ_Bleaching_) or the release from the DNA after termination at time *T*. **c)** Exemplary intensity traces for simulated protein synthesis spots with different bleaching rates τ and a transcription initiation rate of 1/5 min^-1^. **d)** ACFs on a linear and log-scaled x-axis for k_tx_ = 1/5 min^-1^ and indicated bleaching rates and gene lengths. As the bleaching rate becomes the rate-limiting step, the ACFs transition from a linear to an exponential decay. **e)** Same as panel (d) for τ_Bleaching_ = 1.5 min and different transcription initiation rates and gene lengths. Here, the offset at long timescales approaches zero for faster initiation rates. **f)** Autocorrelation functions for the simulated intensity traces and mono-exponential fits (solid lines) for two different bleaching rates (as indicated on the right hand-side of the graph) and three different gene constructs. **g)** The inverse amplitude, G(0)^-1^, and correlation time $\tau_{C}$ for various combinations of transcription elongation rates (v), initiation rates (k_tx_), and bleaching rates (values as in panel a from top to bottom). Each data point is the median of 100 simulated protein synthesis spots.

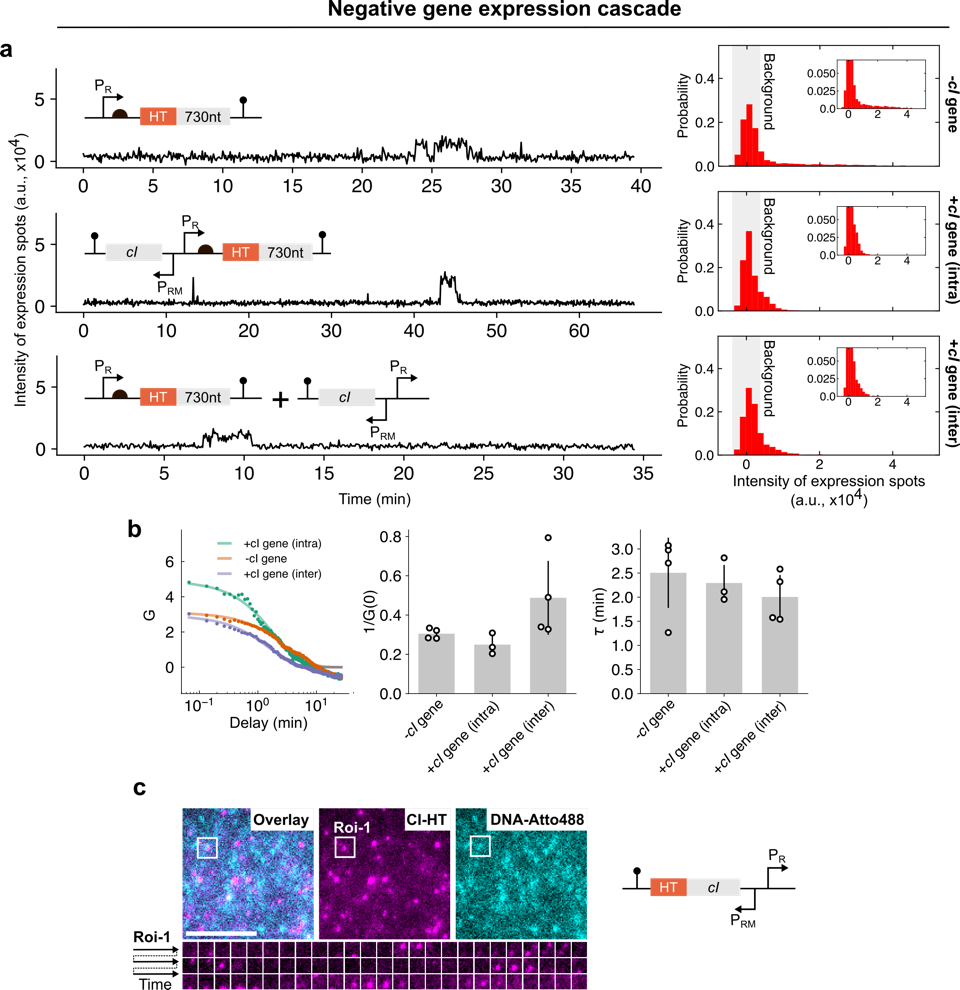

**Supplementary Figure 7: Negative cascaded gene expression reactions on the same DNA molecule. a)** Examples of protein synthesis traces (left) and intensity distributions of expression spots over the ensemble (right) for the three different experiments without *cI* gene (n=3), with *cI* gene encoded on the same DNA molecule as the *ht* gene (intramolecular case, n=3), and the *cI* and *ht* gene split between two DNA molecules (n=4) immobilized in the same flow channel. The *cI* repressor gene and regulatory elements were added to the DNA encoding the HT with 730 nt-long C-terminal extension. **b)** The autocorrelation function (G) with the extracted inverse amplitude and timescale for the three different experiments as indicated in a. **c)** Fluorescence images of DNA and CI-HT spots. Scale bar, 5 µm. White boxes in fluorescent images show the region Roi-1 from which a time series for the signal of protein synthesis (CI-HT) was extracted.

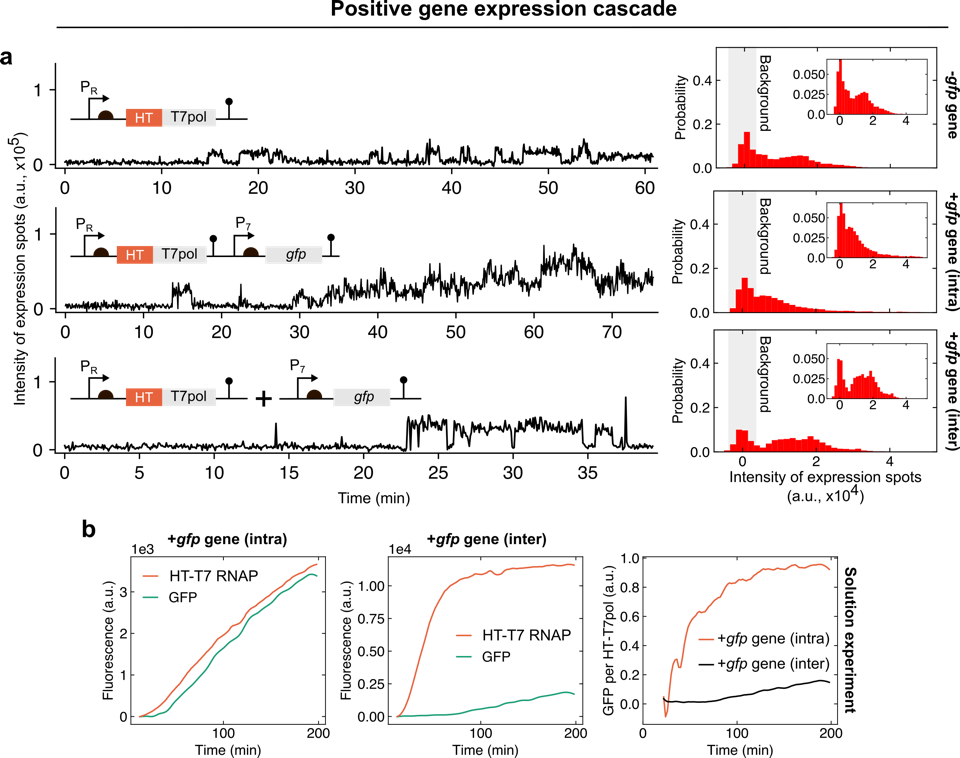

**Supplementary Figure 8: Positive cascaded gene expression reactions on the same DNA molecule. a)** Examples of protein synthesis traces (left) and intensity distributions of expression spots over the ensemble (right) for the three different experimental conditions using DNA molecules encoding the gene for the T7 RNAP (T7 pol) without *gfp* gene (n=5), with encoded T7 RNAP and *gfp* gene (n=4), and the genes for T7 RNAP and GFP split between two DNA molecules (n=3) immobilized in the same flow channel. **b)** Two representative bulk experiments with positive cascade encoded on two different DNA molecules and together on a single DNA molecule (n=2 for each experiment). The rightmost panel shows the GFP signal divided by the HT-T7 RNAP signal.

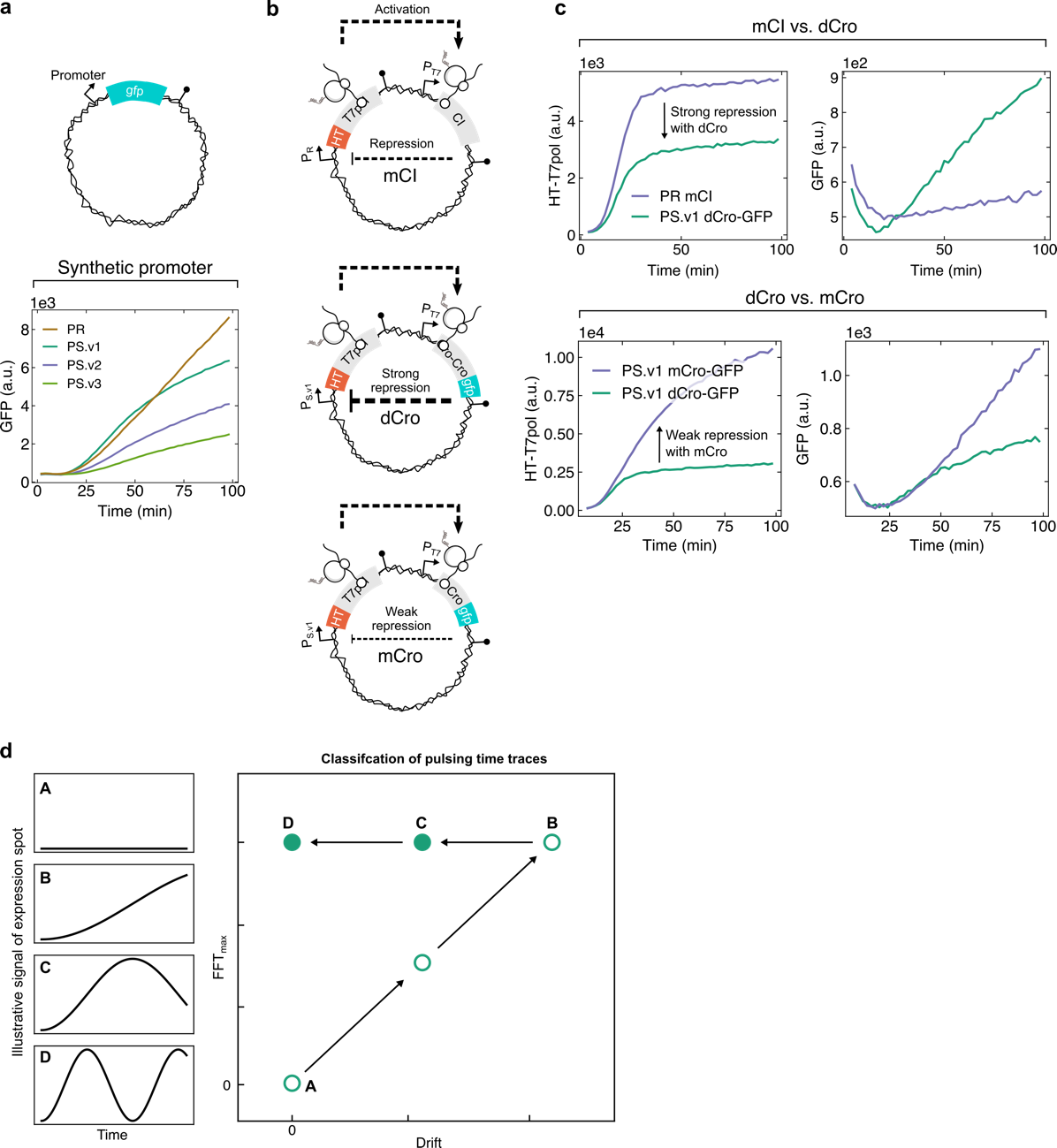

**Supplementary Figure 9: Full circuits with positive and negative feedback on the same DNA molecule. a)** Representative bulk experiment with GFP gene expression for the standard promoter P_R_ and three *de novo* designed promoters (**Supplementary** **Table 1**). Ribosomal binding sites were identical for all the constructs (n=2). **b)** Three different constructs encoding genes for the T7 RNAP and monomeric repressor cI (mCI, left schematic), strong repressor dCro-GFP (dCro, center schematic), and very weak monomeric repressor mCro-GFP (mCro, right schematic). Each promoter controlling the transcription of HT-T7 RNAP contained two strong binding sites for the corresponding repressors (see **Supplementary** **Table 1**). **c)** First two plots show representative bulk expression experiments from mCI and dCro-GFP constructs with HT-T7 RNAP (left panel) and dCro-GFP (right panel) signal. The auto-fluorescence of the cell lysate produced a background signal in the GFP channel for the mCI construct (n=2 for each experiment). The last two plots show representative bulk expressions from dCro-GFP and mCro-GFP constructs with the signals for HT-T7 RNAP and dCro-GFP (or mCro-GFP) signal (n=2 for each experiment). **d)** Illustrative HT-T7 RNAP expression traces with four qualitatively different characteristics (from top to bottom): no activity, accumulation, slow pulse, two pulses. Trace A and B are located along the diagonal on a plot for drift and maximal FFT amplitude ($\mathcal{F}_{max}$). Pulsing is characterized by a constant $\mathcal{F}_{max}$, but reduced drift value (Trace C and D).

**Supplementary Table 1: The DNA sequences with ribosomal binding site, various open reading frames, and terminators.**

| Name | Sequence (RBS is highlighted in red, terminator is underlined in capital letters) |
| --- | --- |
| HaloTag9 gene (capitalized letters are silent mutations) | gaAATAATTTTGTTTAACTTTAAGAAGGAGATATAccatgatcggtactggctttccattcgacccccattatgtggaagtcctgggcgagcgcatgcactacgtcgatgttggtccgcgcgatggcacccctgtgctgttcctgcacggtaacccgacctcctcctacgtgtggcgcaacatcatcccgcatgttgcaccgacccatcgctgcattgctccagacctgatcggtatgggcaaatccgacaaaccagacctgggttatttcttcgacgaccacgtccgcttcatggatgccttcatcgaagccctgggtctggaagaggtcgtcctggtcattcacgactggggctccgctctgggtttccactgggccaagcgcaatccagagcgcgtcaaaggtattgcatttatggagttcatccgccctatcccgacctgggacgaatggccagaatttgcccgcgagaccttccaggccttccgcaccaccgacgtcggccgcaagctgatcatTgatcaCaacgtttttatcgagggtacgctgcGCatgggtgtcgtccgcccgctgactgaagtcgagatggaccattaccgcgagccgttcctgaatcctgttgaccgcgagccactgtggcgcttcccaaacgagctgccaatcgccggtgagccagcgaacatcgtcgcgctggtcgaagaatacatggactggctgcaccagtcccctgtcccgaagctgctgttctggggcaccccaggcgttctgatcccaccggccgaagccgctcgcctggccaaaagcctgcctaactgcaaggctgtggacatcggcccgggtctgaatctgctgcaagaagacaacccggacctgatcggcagcgagatcgcgcgctggctgtctactctggagattgcagcaaacgacgaaaactacgctttagatgactaactcgagCAAAGCCCGCCGAAAGGCGGGCTTTTCTGTgtcg |
| HaloTag9 with truncated gp3 fusion (capitalized letters is the KRAPGTS linker and silent mutations) | gcAATAATTTTGTTTAACTTTAAGAAGGAGATATAccatgatcggtactggctttccattcgacccccattatgtggaagtcctgggcgagcgcatgcactacgtcgatgttggtccgcgcgatggcacccctgtgctgttcctgcacggtaacccgacctcctcctacgtgtggcgcaacatcatcccgcatgttgcaccgacccatcgctgcattgctccagacctgatcggtatgggcaaatccgacaaaccagacctgggttatttcttcgacgaccacgtccgcttcatggatgccttcatcgaagccctgggtctggaagaggtcgtcctggtcattcacgactggggctccgctctgggtttccactgggccaagcgcaatccagagcgcgtcaaaggtattgcatttatggagttcatccgccctatcccgacctgggacgaatggccagaatttgcccgcgagaccttccaggccttccgcaccaccgacgtcggccgcaagctgatcatTgatcaCaacgtttttatcgagggtacgctgcGCatgggtgtcgtccgcccgctgactgaagtcgagatggaccattaccgcgagccgttcctgaatcctgttgaccgcgagccactgtggcgcttcccaaacgagctgccaatcgccggtgagccagcgaacatcgtcgcgctggtcgaagaatacatggactggctgcaccagtcccctgtcccgaagctgctgttctggggcaccccaggcgttctgatcccaccggccgaagccgctcgcctggccaaaagcctgcctaactgcaaggctgtggacatcggcccgggtctgaatctgctgcaagaagacaacccggacctgatcggcagcgagatcgcgcgctggctgtctactctggagattAAGCGAGCTCCCGGGACCAGCtcgcaagctctgcaacaaatttttaaccaagcaaatacaactaactttgtagtatcaataccacatagtaatactacatctgcttttactttaaatgctcagtcagttcctattccaggaattagaatacctgttactgataccgtgactgggccgtttggactgggccgagcacaacgtccaggtgttacatttgagtatgatccactcattgtgagatttatagttgatgaagaacttaagtcgtggataggaatgtatgaatggatgctaggaactagcaactatcttacaggtgaaaatactgcccaaaaaacaggtcctgagtacattacgctttacatcttagataatagcaaaactgaaatcgtgatgtcaataaatttttataagccttgggtttctgacctatctgaagtagaatttagctacacggaagattcagacccggctttagtatgtactgcaacaattccttacacctattttcaagtagaaaaagatggtaaaattatagcagaagttgcagcaaacgacgaaaactacgctttagatgactaactcgagCAAAGCCCGCCGAAAGGCGGGCTTTTCTGTgtcg |
| HaloTag9 with truncated gp48 fusion (capitalized letters is the KRAPGTS linker and silent mutations) | gcAATAATTTTGTTTAACTTTAAGAAGGAGATATAccatgatcggtactggctttccattcgacccccattatgtggaagtcctgggcgagcgcatgcactacgtcgatgttggtccgcgcgatggcacccctgtgctgttcctgcacggtaacccgacctcctcctacgtgtggcgcaacatcatcccgcatgttgcaccgacccatcgctgcattgctccagacctgatcggtatgggcaaatccgacaaaccagacctgggttatttcttcgacgaccacgtccgcttcatggatgccttcatcgaagccctgggtctggaagaggtcgtcctggtcattcacgactggggctccgctctgggtttccactgggccaagcgcaatccagagcgcgtcaaaggtattgcatttatggagttcatccgccctatcccgacctgggacgaatggccagaatttgcccgcgagaccttccaggccttccgcaccaccgacgtcggccgcaagctgatcatTgatcaCaacgtttttatcgagggtacgctgcGCatgggtgtcgtccgcccgctgactgaagtcgagatggaccattaccgcgagccgttcctgaatcctgttgaccgcgagccactgtggcgcttcccaaacgagctgccaatcgccggtgagccagcgaacatcgtcgcgctggtcgaagaatacatggactggctgcaccagtcccctgtcccgaagctgctgttctggggcaccccaggcgttctgatcccaccggccgaagccgctcgcctggccaaaagcctgcctaactgcaaggctgtggacatcggcccgggtctgaatctgctgcaagaagacaacccggacctgatcggcagcgagatcgcgcgctggctgtctactctggagattAAGCGAGCTCCCGGGACCAGCgcaattgttaaagaaataactgctgatttaattaaaaagtccggtgagaaaatttcagccggacagagtactaaatcagaagtaggaactaaaacatacacagcccagtttccaactgggcgtgctagtggtaatgacactacagaggacttccaggtaacagatctatataagaatggattattatttactgcatacaatatgtcatctagggattctggaagtcttagatcgatgagatctaactactcttcttcatcttcgagtattttacgtacagctagaaacactattagtagtacagtatcaaaactatcaaatggattaatatcaaataataattcaggaacaataagtaaatctcctatcgcaaacattcttttaccgagatctaaatctgatgttgatacatcatcacatagatttaatgatgttcaagaaagccttatcagtagaggcggaggtactgctactggtgtgctaagtaatattgcttcaaccgcagtatttggggcactggaaagtataacacaaggtataatggctgataataatgaacagatttatacgacagccagaagtatgtatggtggtgctgaaaatagaactaaagtgtttacatgggatttgactccacgttcaacagaagatttaatggctattattaatatctatcaatattttaactattttcttatggtgaaacgggtaaatctcaatatgctgctgaaataaaggggtatttagatgattggtatcgttctacgttaattgaacctttatctccggaagacgcagctaaaaataaaacactatttgagaaaatgacatcgagtttaactaacgttctagtagtttcaaacccgacagtttggatggtgaaaaactttggcgcaacatctaagtttgatggaaaaacggaaatatttggtccatgtcaaatacagagcattagatttgataaaacacctaatggtaactttaacggattagctattgctccaaacctccctagtacatttactctcgagattactatgagagaaattatcacgttaaaccgtgcttctttatatgcggggacttttgcagcaaacgacgaaaactacgctttagatgactaactcgagCAAAGCCCGCCGAAAGGCGGGCTTTTCTGTgtcg |
| HaloTag9 with gp6 fusion (capitalized letters is the KRAPGTS linker and silent mutations) | gcAATAATTTTGTTTAACTTTAAGAAGGAGATATAccatgatcggtactggctttccattcgacccccattatgtggaagtcctgggcgagcgcatgcactacgtcgatgttggtccgcgcgatggcacccctgtgctgttcctgcacggtaacccgacctcctcctacgtgtggcgcaacatcatcccgcatgttgcaccgacccatcgctgcattgctccagacctgatcggtatgggcaaatccgacaaaccagacctgggttatttcttcgacgaccacgtccgcttcatggatgccttcatcgaagccctgggtctggaagaggtcgtcctggtcattcacgactggggctccgctctgggtttccactgggccaagcgcaatccagagcgcgtcaaaggtattgcatttatggagttcatccgccctatcccgacctgggacgaatggccagaatttgcccgcgagaccttccaggccttccgcaccaccgacgtcggccgcaagctgatcatTgatcaCaacgtttttatcgagggtacgctgcGCatgggtgtcgtccgcccgctgactgaagtcgagatggaccattaccgcgagccgttcctgaatcctgttgaccgcgagccactgtggcgcttcccaaacgagctgccaatcgccggtgagccagcgaacatcgtcgcgctggtcgaagaatacatggactggctgcaccagtcccctgtcccgaagctgctgttctggggcaccccaggcgttctgatcccaccggccgaagccgctcgcctggccaaaagcctgcctaactgcaaggctgtggacatcggcccgggtctgaatctgctgcaagaagacaacccggacctgatcggcagcgagatcgcgcgctggctgtctactctggagattAAGCGAGCTCCCGGGACCAGCgcaaatacccctgtaaattatcaattaacaagaacagcaaatgctattcccgagatattcgtcgggggtacatttgctgaaataaaacaaaacctcattgaatggcttaatggccaaaatgaatttttggattatgattttgaaggctcaagattaaacgttctgtgtgaccttttagcttataatacattatacattcagcagtttggtaatgctgctgtgtatgaaagctttatgcgtactgctaacttacgaagttcagttgttcaagctgcacaagataacggatatttacctacttcaaaatccgctgcgcagaccgaaattatgttaacatgcactgacgcattgaataggaattacattactattcctcgcggaactcgctttttagcatatgcaaaagatacttctgttaatccatataacttcgtttctaccgaagacgttattgctattcgtgataaaaataaccaatattttccgcgtttaaaattggcccagggacgtatagtaagaactgaaatcatttatgataaattaacacctattatcatttatgataaaaatattgatagaaaccaggttaaattatacgttgatggagcggaatggattaactggacgagaaagtcaatggttcatgctggttcaacatcaacgatttactatatgcgtgaaactattgatggaaacactgaattctattttggtgaaggtgaaatttctgttaatgcttctgaaggagctttgaccgctaattatatcggaggtcttaaacctactcagaactctacgattgttattgagtacattagtactaatggtgctgacgcgaacggagcagtcggattttcatacgcagatacattaacaaatataactgtcatcaatattaatgaaaatccaaacgatgatccagattttgttggggcagatggaggcggtgatccagaagatattgagcgtattcgcgaattgggtactattaaacgcgaaacccaacaacgatgcgtaactgcgactgactatgatacattcgtttcagagagatttggttctattattcaagctgttcagactttcactgattctactaaacctgggtatgcatttattgctgctaaacctaaatcaggattgtatttaactaccgtacagcgtgaagatattaaaaattatctcaaagactataatttagctcctattacgccatcaattatttctcctaattatctttttattaagactaatttaaaagtcacatatgctttaaataaactgcaagaatccgaacagtggcttgaaggtcaaataattgataaaatagatcgctattataccgaagatgtagaaatttttaactcgtctttcgctaaatctaagatgttgacatatgtagatgatgcagatcattctgtcattggttcatcagcgactattcaaatggttcgtgaagtacaaaacttctataaaacgcctgaagcgggtattaaatacaataatcaaataaaagatcgttctatggaatctaatacgttttcatttaattctggacgaaaggttgtaaatcctgatactggtttagaagaagatgtattatatgacgttcgtatagtatcaacagaccgagattctaaaggaattggtaaagttattattggtccatttgcttctggcgatgttacagaaaatgaaaacattcagccgtatacaggcaacgattttaacaaattagcaaattctgatggacgcgacaaatactatgttatcggtgaaataaattatccagctgatgtgatttattggaatatcgctaaaattaatttaacatctgaaaaatttgaagttcagaccattgaattatattctgacccaaccgatgatgttatctttactcgcgatggttcactgattgtatttgaaaatgacttacgtccacaatacttaactatcgatttggagcctatatcacaagcagcaaacgacgaaaactacgctttagatgactaactcgagCAAAGCCCGCCGAAAGGCGGGCTTTTCTGTgtcg |
| HaloTag9 with truncated gp7 fusion (capitalized letters is the KRAPGTS linker and silent mutations) | gcAATAATTTTGTTTAACTTTAAGAAGGAGATATAccatgatcggtactggctttccattcgacccccattatgtggaagtcctgggcgagcgcatgcactacgtcgatgttggtccgcgcgatggcacccctgtgctgttcctgcacggtaacccgacctcctcctacgtgtggcgcaacatcatcccgcatgttgcaccgacccatcgctgcattgctccagacctgatcggtatgggcaaatccgacaaaccagacctgggttatttcttcgacgaccacgtccgcttcatggatgccttcatcgaagccctgggtctggaagaggtcgtcctggtcattcacgactggggctccgctctgggtttccactgggccaagcgcaatccagagcgcgtcaaaggtattgcatttatggagttcatccgccctatcccgacctgggacgaatggccagaatttgcccgcgagaccttccaggccttccgcaccaccgacgtcggccgcaagctgatcatTgatcaCaacgtttttatcgagggtacgctgcGCatgggtgtcgtccgcccgctgactgaagtcgagatggaccattaccgcgagccgttcctgaatcctgttgaccgcgagccactgtggcgcttcccaaacgagctgccaatcgccggtgagccagcgaacatcgtcgcgctggtcgaagaatacatggactggctgcaccagtcccctgtcccgaagctgctgttctggggcaccccaggcgttctgatcccaccggccgaagccgctcgcctggccaaaagcctgcctaactgcaaggctgtggacatcggcccgggtctgaatctgctgcaagaagacaacccggacctgatcggcagcgagatcgcgcgctggctgtctactctggagattAAGCGAGCTCCCGGGACCAGCatgacagtaaaagcaccttcagtcactagtctcagaatttccaagttatccgcaaatcaggtgcaagtacgctgggatgacgttggtgctaatttctactattttgtagaaatcgctgagacaaaaacaaactcgggggaaaatctcccgagtaatcaatatcgttggattaatttaggatatacagcaaataatagtttcttttttgatgatgctgatccattaacaacatacattattagagtagccacagctgcgcaagattttgagcagtctgattggatttataccgaagagtttgaaacttttgctacaaatgcttatacatttcaaaacatgattgaaatgcaattagccaataaattcattcaggaaaaatttactcttaataattctgattatgttaattttaataatgatactataatggctgcattgatgaatgaatcattccaattcagcccatcgtatgttgatgtttcatcaataagtaattttattattggtgaaaatgagtatcatgaaatacaaggttctattcagcaagtatgtaaggatattaaccgagtttatttgatggaatcagaaggaattctatatctttttgagcgctatcaacctgtagttaaagtatccaatgataaaggacaaacctggaaagctgtaaagctcttcaatgaccgtgtaggatatcctttatctaagacagtatattaccaatctgcgaacacaacatacgttctaggatacgacaagattttctatggccgcaaatctactgatgttagatggtcagccgatgatgtcagatttagttctcaggatataacatttgctaaacttggcgaccaattacatctaggatttgatgtagaaatttttgccacttacgcgactttaccagcgaatgtataccgcattgcagaagctattacttgcaccgatgattacatttacgttgtcgccagagacaaagttagatacataaaaacgagtaatgcacttatagattttgatccattatctccaacatattcggaaagactttttgaacctgataccatgactataaccggaaatcctaaagcagtatgctataaaatggattctatctgtgataaagtttttgctcttattattggtgaagttgaaacattaaatgctaatcctagaacatcaaaaataattgattccgctgataaaggaatatatgttttaaatcatgacgaaaaaacatggaaaagagtttttggtaataccgaagaagaaagaagacgtattcaacccggatatgcgaatatgtcaactgacggtaaattagtttctctgtcttcgagtaattttaaatttttaagtgataatgttgttaatgaccctgaaactgcagcaaaatatcagttaattggcgctgttaaatatgaatttcctcgtgaatggttagctgataagcattatcatatgatggcatttatagcggatgaaacatctgattgggagacttttactcctcaaccaatgaaatactacgcagaaccattctttaactggtctaaaaaatctaacacacgttgttggataaacaactctgatagagctgtggtagtttatgctgatttaaaatacactaaagttatagaaaatattccggaaacatcaccagatagattagttcatgaatactgggatgatggtgattgcactatagtaatgccaaatgtcaaattcactggatttaaaaaatacgcatcaggaatgcttttctataaagcctccggtgaaataatttcttactatgattttaactatcgtgtgagagatacagtagaaattatttggaagccaactgaagtatttttaaaagcatttttacaaaaccaagagcatgagactccttggtcaccagaagaagagcgtggattagctgaccctgatttaagaccattaattggcacaatgatgcctgattcttatttgttacaggattcgaattttgaggcattttgcgaagcatatattcagtatctttctgcagcaaacgacgaaaactacgctttagatgactaactcgagCAAAGCCCGCCGAAAGGCGGGCTTTTCTGTgtcgacc |
| mvenus gene with truncated gp48 fusion (capitalized letters is the KRAPGTS linker and silent mutations) | gcAATAATTTTGTTTAACTTTAAGAAGGAGATATAccatggagcttttcactggcgttgttcccatcctggtcgagctggacggcgacgtaaacggccacaagttcagcgtgtccggcgagggcgagggcgatgccacctacggcaagctgaccctgaagctgatctgcaccaccggcaagctgcccgtgccctggcccaccctcgtgaccaccctgggctacggcgttcagtgcttcgcccgctaccccgaccacatgaagcagcacgacttcttcaagtccgccatgcccgaaggctacgtccaggagcgcaccatcttcttcaaggacgacggcaactacaagacccgcgccgaggtgaagttcgagggcgacaccctggtgaaccgcatcgagctgaagggcatcgacttcaaggaggacggcaacatcctggggcacaagctggagtacaactacaacagccacaacgtctatatcaccgccgacaagcagaagaacggcatcaaggccaacttcaagatccgccacaacatcgaggacggcggcgtgcagctcgccgaccactaccagcagaacacccccatcggcgacggccccgtgctgctgcccgacaaccactacctgagctaccagtccaagctgagcaaagaccccaacgagaagcgcgatcacatggtcctgctggagttcgtgaccgccgccgggatcAAGCGAGCTCCCGGGACCAGCgcaattgttaaagaaataactgctgatttaattaaaaagtccggtgagaaaatttcagccggacagagtactaaatcagaagtaggaactaaaacatacacagcccagtttccaactgggcgtgctagtggtaatgacactacagaggacttccaggtaacagatctatataagaatggattattatttactgcatacaatatgtcatctagggattctggaagtcttagatcgatgagatctaactactcttcttcatcttcgagtattttacgtacagctagaaacactattagtagtacagtatcaaaactatcaaatggattaatatcaaataataattcaggaacaataagtaaatctcctatcgcaaacattcttttaccgagatctaaatctgatgttgatacatcatcacatagatttaatgatgttcaagaaagccttatcagtagaggcggaggtactgctactggtgtgctaagtaatattgcttcaaccgcagtatttggggcactggaaagtataacacaaggtataatggctgataataatgaacagatttatacgacagccagaagtatgtatggtggtgctgaaaatagaactaaagtgtttacatgggatttgactccacgttcaacagaagatttaatggctattattaatatctatcaatattttaactattttcttatggtgaaacgggtaaatctcaatatgctgctgaaataaaggggtatttagatgattggtatcgttctacgttaattgaacctttatctccggaagacgcagctaaaaataaaacactatttgagaaaatgacatcgagtttaactaacgttctagtagtttcaaacccgacagtttggatggtgaaaaactttggcgcaacatctaagtttgatggaaaaacggaaatatttggtccatgtcaaatacagagcattagatttgataaaacacctaatggtaactttaacggattagctattgctccaaacctccctagtacatttactctcgagattactatgagagaaattatcacgttaaaccgtgcttctttatatgcggggacttttgcagcaaacgacgaaaactacgctttagatgactaactcgagCAAAGCCCGCCGAAAGGCGGGCTTTTCTGTgtcgacc |
| N-terminal HA tag for HT (HA tag highlighted with bold letters) until start codon “atg” | gcAATAATTTTGTTTAACTTTAAGAAGGAGATATAcc**ATGACCAGCTACCCATACGATGTTCCAGATTACGCTGGCCGCTTAATTAAACATATGACC**atg… |
| P_S.V1_ (Cro binding sites are underlined) until start codon “atg” | gttccgctgggcattctatcaccgcgggtgataaacttgacaactatcccttgcggtgatattatggcctgctatgcagctagcAATAATTTTGTTTAACTTTAAGAAGGAGATATAccatg… |
| P_S.V2_ (Cro binding sites are underlined) until start codon “atg” | gttccgctgggcattctatcaccgcgggtgataaaattgacatttatcccttgcggtgatataatgctatgctatgcagctagcAATAATTTTGTTTAACTTTAAGAAGGAGATATAccatg… |
| P_S.V3_ (strong CI, but weak Cro binding sites are underlined) until start codon “atg” | gttccgctgggcattaacaccgtgcgtgttgtattgacactacctctggcggtgatattattatctgctatgcagctagcAATAATTTTGTTTAACTTTAAGAAGGAGATATAccatg… |
| DNA molecule encoding the full circuit with HT-T7 RNAP transcribed under the PS.V1 promoter (underlined, lowercase letters) and dCro-GFP transcribed under the T7 promoter (underlined, lowercase letters) | cgccgcagagtggatgtttgacatggtgaagactatcgcaccatcagccagaaaaccgaattttgctgggtgggctaacgatatccgcctgatgcgtgaacgtgacggacgtaaccaccgcgacatgtgtgtgctgttccgctgggcattctatcaccgcgggtgataaacttgacaactatcccttgcggtgatattatggcctgctatgcagctagcAATAATTTTGTTTAACTTTAAGAAGGAGATATAccatgatcggtactggctttccattcgacccccattatgtggaagtcctgggcgagcgcatgcactacgtcgatgttggtccgcgcgatggcacccctgtgctgttcctgcacggtaacccgacctcctcctacgtgtggcgcaacatcatcccgcatgttgcaccgacccatcgctgcattgctccagacctgatcggtatgggcaaatccgacaaaccagacctgggttatttcttcgacgaccacgtccgcttcatggatgccttcatcgaagccctgggtctggaagaggtcgtcctggtcattcacgactggggctccgctctgggtttccactgggccaagcgcaatccagagcgcgtcaaaggtattgcatttatggagttcatccgccctatcccgacctgggacgaatggccagaatttgcccgcgagaccttccaggccttccgcaccaccgacgtcggccgcaagctgatcatTgatcaCaacgtttttatcgagggtacgctgcGCatgggtgtcgtccgcccgctgactgaagtcgagatggaccattaccgcgagccgttcctgaatcctgttgaccgcgagccactgtggcgcttcccaaacgagctgccaatcgccggtgagccagcgaacatcgtcgcgctggtcgaagaatacatggactggctgcaccagtcccctgtcccgaagctgctgttctggggcaccccaggcgttctgatcccaccggccgaagccgctcgcctggccaaaagcctgcctaactgcaaggctgtggacatcggcccgggtctgaatctgctgcaagaagacaacccggacctgatcggcagcgagatcgcgcgctggctgtctactctggagattGGTGGTTCCGGTGGTatgaacacgattaacatcgctaagaacgacttctctgacatcgaactggctgctatcccgttcaacactctggctgaccattacggtgagcgtttagctcgcgaacagttggcccttgagcatgagtcttacgagatgggtgaagcacgcttccgcaagatgtttgagcgtcaacttaaagctggtgaggttgcggataacgctgccgccaagcctctcatcactaccctactccctaagatgattgcacgcatcaacgactggtttgaggaagtgaaagctaagcgcggcaagcgcccgacagccttccagttcctgcaagaaatcaagccggaagccgtagcgtacatcaccattaagaccactctggcttgcctaaccagtgctgacaatacaaccgttcaggctgtagcaagcgcaatcggtcgggccattgaggacgaggctcgcttcggtcgtatccgtgaccttgaagctaagcacttcaagaaaaacgttgaggaacaactcaacaagcgcgtagggcacgtctacaagaaagcatttatgcaagttgtcgaggctgacatgctctctaagggtctactcggtggcgaggcgtggtcttcgtggcataaggaagactctattcatgtaggagtacgctgcatcgagatgcCcattgagtcaaccggaatggttagcttacaccgccaaaatgctggcgtagtaggtcaagactctgagactatcgaactcgcacctgaatacgcGgaggctatcgcaacccgtgcaggtgcgctggctggcatctctccgatgttccaaccttgcgtagttcctcctaagccgtggactggcattactggtggtggctattgggctaacggtcgtcgtcctctggcgctggtgcgtactcacagtaagaaagcactgatgcgctacgaagacgtttacatgcctgaggtgtacaaagcgattaacattgcgcaaaacaccgcatggaaaatcaacaagaaagtcctagcggtcgccaacgtaatcaccaagtggaagcattgtccggtcgaggacatccctgcgattgagcgtgaagaactcccgatgaaaccggaagacatcgacatgaatcctgaggctctcaccgcgtggaaacgtgctgccgctgctgtgtaccgcaaggacaaggctcgcaagtctcgccgtatcagccttgagttcatgcttgagcaagccaataagtttgctaaccataaggccatctggttcccttacaacatggactggcgcggtcgtgtttacgctgtgtcaatgttcaacccgcaaggtaacgatatgaccaaaggactgcttacgctggcgaaaggtaaaccaatcggtaaggaaggttactactggctgaaaatccacggtgcaaactgtgcgggtgtcgataaggttccgttccctgagcgcatcaagttcattgaggaaaaccacgagaacatcatggcttgcgctaagtctccactggagaacacttggtgggctgagcaagattctccgttctgcttccttgcgttctgctttgagtacgctggggtacagcaccacggcctgagctataactgctcccttccgctggcgtttgacgggtcttgctctggcatccagcacttctccgcgatgctccgagatgaggtaggtggtcgcgcggttaacttgcttcctagtgaaaccgttcaggacatctacgggattgttgctaagaaagtcaacgagattctacaagcagacgcaatcaatgggaccgataacgaagtagttaccgtgaccgatgagaacactggtgaaatctctgagaaagtcaagctgggcactaaggcactggctggtcaatggctggcttacggtgttactcgcagtgtgactaagcgttcagtcatgacgctggcttacgggtccaaagagttcggcttccgtcaacaagtgctggaagataccattcagccagctattgattccggcaagggtctgatgttcactcagccgaatcaggctgctggatacatggctaagctgatttgggaatctgtgagcgtgacggtggtagctgcggttgaagcaatgaactggcttaagtctgctgctaagctgctggctgcGgaggtcaaagataagaagactggagagattcttcgcaagcgttgcgctgtgcattgggtaactcctAatggtttccctgtgtggcaggaatacaagaagcctattcagacgcgcttgaacctgatgttcctcggtcagttccgcttacagcctaccattaacaccaacaaagatagcgagattgatgcacacaaacaggagtctggtatcgctcctaactttgtacacagccaagacggtagccaccttcgtaagactgtagtgtgggcacacgagaagtacggaatcgaatcttttgcactgattcacgactccttcggtaccattccggctgacgctgcgaacctgttcaaagcagtgcgcgaaactatggttgacacatatgagtcttgtgatgtactggctgatttctacgaccagttcgctgaccagttgcacgagtctcaattggacaaaatgccagcacttccggctaaaggtaacttgaacctccgtgacatcttagagtcggacttcgcgttcgcgtaactcgagCAAAGCCCGCCGAAAGGCGGGCTTTTCTGTgtcgaccgatgcccttgagagccttcaacccagtcagctccttccggtgggcgcggggcatgactatcgtcgccgcacttatgactgtcttctttatcatgcaactcgtaggacaggtgccggcagcgctcttccgaggctattcggctatgactgggcataatacgactcactataggggttgcagctagcAATAATTTTGTTTAACTTTAAGAAGGAGGTATAcatatggagcagcgtatcacattaaaagactacgctatgcgtttcgggcagacaaagactgcaaaagacctgggagtatatcagtccgcgattaacaaggcgattcatgcgggacgtaaaatcttcttaaccatcaacgcagacgggtcagtttatgcggaagaggtaaaaccatttccgtccaataaaaaaactaccgcagcagcaggtggaggtacaggaggtggatccatggaacagcgcattaccctgaaagattatgcaatgcgttttggacagaccaaaaccgcgaaagatctgggtgtgtaccaatcggcgatcaataaggccattcatgcaggccgcaagatttttttaactatcaacgctgacggcagcgtttatgcggaggaagtcaaacctttcccgtcaaacaaaaaaacgacagccagaagagcaaagaaggaagccatggagcttttcactggcgttgttcccatcctggtcgagctggacggcgacgtaaacggccacaagttcagcgtgtccggcgagggcgagggcgatgccacctacggcaagctgaccctgaagttcatctgcaccaccggcaagctgcccgtgccctggcccaccctcgtgaccaccctgacctacggcgtgcagtgcttcagccgctaccccgaccacatgaagcagcacgacttcttcaagtccgccatgcccgaaggctacgtccaggagcgcaccatcttcttcaaggacgacggcaactacaagacccgcgccgaggtgaagttcgagggcgacaccctggtgaaccacatcgagctgaagggcatcgacttcaaggaggacggcaacatcctggggcacaagctggagtacaactacaacagccacaacgtctatatcatggccgacaagcagaagaacggcatcaaggtgaacttcaagatccgccacaacatcgaggacggcagcgtgcagctcgccgaccactaccagcagaacacccccatcggcgacggccccgtgctgctgcccgacaaccactacctgagcacccagtccgccctgagcaaagaccacaacgagaagcgcgatcacatggtcctgctggagttcgtgaccgccgccgggatcgcagcaaacgacgaaaactacgctttagatgactaactcgagCAAAGCCCGCCGAAAGGCGGGCTTTTCTGTgtcgaccgatgcccttgagagccttcaacccagtcagctccttccggtgggcgcggggcatgactatcgtcgccgcacttatgactgtcttctttatcatgcaactcgtaggacaggcaacagacaatcggctgctctgatgctgcgctcggtcgttcggctgcggcgagcggtatcagctcactcaaaggcggtaatacggttatccacagaatcaggggataacgcaggaaagaacatgtgagcaaaaggccagcaaaaggccaggaaccgtaaaaaggccgcgttgctggcgtttttccataggctccgcccccctgacgagcatcacaaaaatcgacgctcaagtcagaggtggcgaaacccgacaggactataaagataccaggcgtttccccctggaagctccctcgtgcgctctcctgttccgaccctgccgcttaccggatacctgtccgcctttctcccttcgggaagcgtggcgctttctcaatgctcacgctgtaggtatctcagttcggtgtaggtcgttcgctccaagctgggctgtgtgcacgaaccccccgttcagcccgaccgctgcgccttatccggtaactatcgtcttgagtccaacccggtaagacacgacttatcgccactggcagcagccactggtaacaggattagcagagcgaggtatgtaggcggtgctacagagttcttgaagtggtggcctaactacggctacactagaaggacagtatttggtatctgcgctctgctgaagccagttaccttcggaaaaagagttggtagctcttgatccggcaaacaaaccaccgctggtagcggtggtttttttgtttgcaagcagcagattacgcgcagaaaaaaaggatctcaagaagatcctttgatcttttctacggggtctgacgctcagtggaacgaaaactcacgttaatcgattttggtcatgagctgaggaaaaaggatcttcacctagatccttttaaattaaaaatgaagttttaaatcaatctaaagtatatatgagtaaacttggtctgacagttaccaatgcttaatcagtgaggcacctatctcagcgatctgtctatttcgttcatccatagttgcctgactccccgtcgtgtagataactacgatacgggagggcttaccatctggccccagtgctgcaatgataccgcgagacccacgctcaccggctccagatttatcagcaataaaccagccagccggaagggccgagcgcagaagtggtcctgcaactttatccgcctccatccagtctattaattgttgccgggaagctagagtaagtagttcgccagttaatagtttgcgcaacgttgttgccattgctacaggcatcgtggtgtcacgctcgtcgtttggtatggcttcattcagctccggttcccaacgatcaaggcgagttacatgatcccccatgttgtgcaaaaaagcggttagctccttcggtcctccgatcgttgtcagaagtaagttggccgcagtgttatcactcatggttatggcagcactgcataattctcttactgtcatgccatccgtaagatgcttttctgtgactggtgagtactcaaccaagtcattctgagaatagtgtatgcggcgaccgagttgctcttgcccggcgtcaatacgggataataccgcgccacatagcagaactttaaaagtgctcatcattggaaaacgttcttcggggcgaaaactctcaaggatcttaccgctgttgagatccagttcgatgtaacccactcgtgcacccaactgatcttcagcatcttttactttcaccagcgtttctgggtgagcaaaaacaggaaggcaaaatgccgcaaaaaagggaataagggcgacacggaaatgttgaatactcatactcttcctttttcaatattattgaagcatttatcagggttattgtctcatgagcggatacatatttgaatgtatttagaaaaataaacaaataggggttccgcgcacatttccccgaaaagtgccacctgacgtctaagaaaccattattatcatgacattaacctataaaaataggcgtatcacgaggccctttcgtcttcaagaattctggcgaatcctctgaccagccagaaaacgacctttctgtggtgaaaccggatgctgcaattcagagcggcagcaagtgggggacagcagaagacctgac |
| DNA molecule encoding the full circuit with HT-T7 RNAP transcribed under the PS.V1 promoter (underlined, lowercase letters) and mCro-GFP transcribed under the T7 promoter (underlined, lowercase letters) | cgccgcagagtggatgtttgacatggtgaagactatcgcaccatcagccagaaaaccgaattttgctgggtgggctaacgatatccgcctgatgcgtgaacgtgacggacgtaaccaccgcgacatgtgtgtgctgttccgctgggcattctatcaccgcgggtgataaacttgacaactatcccttgcggtgatattatggcctgctatgcagctagcAATAATTTTGTTTAACTTTAAGAAGGAGATATAccatgatcggtactggctttccattcgacccccattatgtggaagtcctgggcgagcgcatgcactacgtcgatgttggtccgcgcgatggcacccctgtgctgttcctgcacggtaacccgacctcctcctacgtgtggcgcaacatcatcccgcatgttgcaccgacccatcgctgcattgctccagacctgatcggtatgggcaaatccgacaaaccagacctgggttatttcttcgacgaccacgtccgcttcatggatgccttcatcgaagccctgggtctggaagaggtcgtcctggtcattcacgactggggctccgctctgggtttccactgggccaagcgcaatccagagcgcgtcaaaggtattgcatttatggagttcatccgccctatcccgacctgggacgaatggccagaatttgcccgcgagaccttccaggccttccgcaccaccgacgtcggccgcaagctgatcatTgatcaCaacgtttttatcgagggtacgctgcGCatgggtgtcgtccgcccgctgactgaagtcgagatggaccattaccgcgagccgttcctgaatcctgttgaccgcgagccactgtggcgcttcccaaacgagctgccaatcgccggtgagccagcgaacatcgtcgcgctggtcgaagaatacatggactggctgcaccagtcccctgtcccgaagctgctgttctggggcaccccaggcgttctgatcccaccggccgaagccgctcgcctggccaaaagcctgcctaactgcaaggctgtggacatcggcccgggtctgaatctgctgcaagaagacaacccggacctgatcggcagcgagatcgcgcgctggctgtctactctggagattGGTGGTTCCGGTGGTatgaacacgattaacatcgctaagaacgacttctctgacatcgaactggctgctatcccgttcaacactctggctgaccattacggtgagcgtttagctcgcgaacagttggcccttgagcatgagtcttacgagatgggtgaagcacgcttccgcaagatgtttgagcgtcaacttaaagctggtgaggttgcggataacgctgccgccaagcctctcatcactaccctactccctaagatgattgcacgcatcaacgactggtttgaggaagtgaaagctaagcgcggcaagcgcccgacagccttccagttcctgcaagaaatcaagccggaagccgtagcgtacatcaccattaagaccactctggcttgcctaaccagtgctgacaatacaaccgttcaggctgtagcaagcgcaatcggtcgggccattgaggacgaggctcgcttcggtcgtatccgtgaccttgaagctaagcacttcaagaaaaacgttgaggaacaactcaacaagcgcgtagggcacgtctacaagaaagcatttatgcaagttgtcgaggctgacatgctctctaagggtctactcggtggcgaggcgtggtcttcgtggcataaggaagactctattcatgtaggagtacgctgcatcgagatgcCcattgagtcaaccggaatggttagcttacaccgccaaaatgctggcgtagtaggtcaagactctgagactatcgaactcgcacctgaatacgcGgaggctatcgcaacccgtgcaggtgcgctggctggcatctctccgatgttccaaccttgcgtagttcctcctaagccgtggactggcattactggtggtggctattgggctaacggtcgtcgtcctctggcgctggtgcgtactcacagtaagaaagcactgatgcgctacgaagacgtttacatgcctgaggtgtacaaagcgattaacattgcgcaaaacaccgcatggaaaatcaacaagaaagtcctagcggtcgccaacgtaatcaccaagtggaagcattgtccggtcgaggacatccctgcgattgagcgtgaagaactcccgatgaaaccggaagacatcgacatgaatcctgaggctctcaccgcgtggaaacgtgctgccgctgctgtgtaccgcaaggacaaggctcgcaagtctcgccgtatcagccttgagttcatgcttgagcaagccaataagtttgctaaccataaggccatctggttcccttacaacatggactggcgcggtcgtgtttacgctgtgtcaatgttcaacccgcaaggtaacgatatgaccaaaggactgcttacgctggcgaaaggtaaaccaatcggtaaggaaggttactactggctgaaaatccacggtgcaaactgtgcgggtgtcgataaggttccgttccctgagcgcatcaagttcattgaggaaaaccacgagaacatcatggcttgcgctaagtctccactggagaacacttggtgggctgagcaagattctccgttctgcttccttgcgttctgctttgagtacgctggggtacagcaccacggcctgagctataactgctcccttccgctggcgtttgacgggtcttgctctggcatccagcacttctccgcgatgctccgagatgaggtaggtggtcgcgcggttaacttgcttcctagtgaaaccgttcaggacatctacgggattgttgctaagaaagtcaacgagattctacaagcagacgcaatcaatgggaccgataacgaagtagttaccgtgaccgatgagaacactggtgaaatctctgagaaagtcaagctgggcactaaggcactggctggtcaatggctggcttacggtgttactcgcagtgtgactaagcgttcagtcatgacgctggcttacgggtccaaagagttcggcttccgtcaacaagtgctggaagataccattcagccagctattgattccggcaagggtctgatgttcactcagccgaatcaggctgctggatacatggctaagctgatttgggaatctgtgagcgtgacggtggtagctgcggttgaagcaatgaactggcttaagtctgctgctaagctgctggctgcGgaggtcaaagataagaagactggagagattcttcgcaagcgttgcgctgtgcattgggtaactcctAatggtttccctgtgtggcaggaatacaagaagcctattcagacgcgcttgaacctgatgttcctcggtcagttccgcttacagcctaccattaacaccaacaaagatagcgagattgatgcacacaaacaggagtctggtatcgctcctaactttgtacacagccaagacggtagccaccttcgtaagactgtagtgtgggcacacgagaagtacggaatcgaatcttttgcactgattcacgactccttcggtaccattccggctgacgctgcgaacctgttcaaagcagtgcgcgaaactatggttgacacatatgagtcttgtgatgtactggctgatttctacgaccagttcgctgaccagttgcacgagtctcaattggacaaaatgccagcacttccggctaaaggtaacttgaacctccgtgacatcttagagtcggacttcgcgttcgcgtaactcgagCAAAGCCCGCCGAAAGGCGGGCTTTTCTGTgtcgaccgatgcccttgagagccttcaacccagtcagctccttccggtgggcgcggggcatgactatcgtcgccgcacttatgactgtcttctttatcatgcaactcgtaggacaggtgccggcagcgctcttccgaggctattcggctatgactgggcataatacgactcactataggggttgcagctagcAATAATTTTGTTTAACTTTAAGAAGGAGGTATAcatatggagcagcgtatcacaggaggtggatccatggaacagcgcattaccctgaaagattatgcaatgcgttttggacagaccaaaaccgcgaaagatctgggtgtgtaccaatcggcgatcaataaggccattcatgcaggccgcaagatttttttaactatcaacgctgacggcagcgtttatgcggaggaagtcaaacctttcccgtcaaacaaaaaaacgacagccagaagagcaaagaaggaagccatggagcttttcactggcgttgttcccatcctggtcgagctggacggcgacgtaaacggccacaagttcagcgtgtccggcgagggcgagggcgatgccacctacggcaagctgaccctgaagttcatctgcaccaccggcaagctgcccgtgccctggcccaccctcgtgaccaccctgacctacggcgtgcagtgcttcagccgctaccccgaccacatgaagcagcacgacttcttcaagtccgccatgcccgaaggctacgtccaggagcgcaccatcttcttcaaggacgacggcaactacaagacccgcgccgaggtgaagttcgagggcgacaccctggtgaaccacatcgagctgaagggcatcgacttcaaggaggacggcaacatcctggggcacaagctggagtacaactacaacagccacaacgtctatatcatggccgacaagcagaagaacggcatcaaggtgaacttcaagatccgccacaacatcgaggacggcagcgtgcagctcgccgaccactaccagcagaacacccccatcggcgacggccccgtgctgctgcccgacaaccactacctgagcacccagtccgccctgagcaaagaccacaacgagaagcgcgatcacatggtcctgctggagttcgtgaccgccgccgggatcgcagcaaacgacgaaaactacgctttagatgactaactcgagCAAAGCCCGCCGAAAGGCGGGCTTTTCTGTgtcgaccgatgcccttgagagccttcaacccagtcagctccttccggtgggcgcggggcatgactatcgtcgccgcacttatgactgtcttctttatcatgcaactcgtaggacaggcaacagacaatcggctgctctgatgctgcgctcggtcgttcggctgcggcgagcggtatcagctcactcaaaggcggtaatacggttatccacagaatcaggggataacgcaggaaagaacatgtgagcaaaaggccagcaaaaggccaggaaccgtaaaaaggccgcgttgctggcgtttttccataggctccgcccccctgacgagcatcacaaaaatcgacgctcaagtcagaggtggcgaaacccgacaggactataaagataccaggcgtttccccctggaagctccctcgtgcgctctcctgttccgaccctgccgcttaccggatacctgtccgcctttctcccttcgggaagcgtggcgctttctcaatgctcacgctgtaggtatctcagttcggtgtaggtcgttcgctccaagctgggctgtgtgcacgaaccccccgttcagcccgaccgctgcgccttatccggtaactatcgtcttgagtccaacccggtaagacacgacttatcgccactggcagcagccactggtaacaggattagcagagcgaggtatgtaggcggtgctacagagttcttgaagtggtggcctaactacggctacactagaaggacagtatttggtatctgcgctctgctgaagccagttaccttcggaaaaagagttggtagctcttgatccggcaaacaaaccaccgctggtagcggtggtttttttgtttgcaagcagcagattacgcgcagaaaaaaaggatctcaagaagatcctttgatcttttctacggggtctgacgctcagtggaacgaaaactcacgttaatcgattttggtcatgagctgaggaaaaaggatcttcacctagatccttttaaattaaaaatgaagttttaaatcaatctaaagtatatatgagtaaacttggtctgacagttaccaatgcttaatcagtgaggcacctatctcagcgatctgtctatttcgttcatccatagttgcctgactccccgtcgtgtagataactacgatacgggagggcttaccatctggccccagtgctgcaatgataccgcgagacccacgctcaccggctccagatttatcagcaataaaccagccagccggaagggccgagcgcagaagtggtcctgcaactttatccgcctccatccagtctattaattgttgccgggaagctagagtaagtagttcgccagttaatagtttgcgcaacgttgttgccattgctacaggcatcgtggtgtcacgctcgtcgtttggtatggcttcattcagctccggttcccaacgatcaaggcgagttacatgatcccccatgttgtgcaaaaaagcggttagctccttcggtcctccgatcgttgtcagaagtaagttggccgcagtgttatcactcatggttatggcagcactgcataattctcttactgtcatgccatccgtaagatgcttttctgtgactggtgagtactcaaccaagtcattctgagaatagtgtatgcggcgaccgagttgctcttgcccggcgtcaatacgggataataccgcgccacatagcagaactttaaaagtgctcatcattggaaaacgttcttcggggcgaaaactctcaaggatcttaccgctgttgagatccagttcgatgtaacccactcgtgcacccaactgatcttcagcatcttttactttcaccagcgtttctgggtgagcaaaaacaggaaggcaaaatgccgcaaaaaagggaataagggcgacacggaaatgttgaatactcatactcttcctttttcaatattattgaagcatttatcagggttattgtctcatgagcggatacatatttgaatgtatttagaaaaataaacaaataggggttccgcgcacatttccccgaaaagtgccacctgacgtctaagaaaccattattatcatgacattaacctataaaaataggcgtatcacgaggccctttcgtcttcaagaattctggcgaatcctctgaccagccagaaaacgacctttctgtggtgaaaccggatgctgcaattcagagcggcagcaagtgggggacagcagaagacctgac |
